# Low-frequency phase temporally coordinates multiple working memory operations

**DOI:** 10.64898/2026.04.30.721949

**Authors:** Yun Ding, Paul Cavanah, Ian Fiebelkorn

## Abstract

Working memory unfolds over time, yet how different working memory operations are temporally coordinated remains unclear. Building on prior links between low-frequency neural oscillations and working memory maintenance and retrieval, as well as evidence that low-frequency oscillations help coordinate cognitive functions, we tested whether low-frequency neural oscillations bridge and/or differentiate distinct working memory operations. Specifically, we tested whether low-frequency phase was linked to memory accuracy and event-related neural responses across three operations: (i) encoding, (ii) retrieval, and (iii) distractor processing during maintenance. Using EEG in human participants, we found that encoding and retrieval were most strongly linked to memory accuracy through theta phase (∼4–7 Hz), measured just prior to each task event. Pre-encoding theta phase also modulated the neural response to memory item onset, suggesting that theta phase influences encoding strength. Critically, the theta phase associated with better memory accuracy differed significantly between encoding and retrieval, consistent with temporally distinct and functionally specific states supporting each working memory operation. In contrast, the influence of distractors on memory accuracy was linked to alpha phase (∼8–10 Hz), with distractor occurrence also appearing to re-engage theta-dependent processes associated with encoding and retrieval. Together, these findings suggest that low-frequency neural oscillations provide a temporal framework that bridges multiple operations of working memory.

**Significance:** Working memory (WM) relies on multiple operations that must be coordinated over time, yet how these processes are temporally organized remains unclear. Neural oscillations have been proposed as a timing mechanism for cognition, yet evidence linking distinct oscillatory phases to distinct WM operations remains limited. Here, we show that memory accuracy depends on the phase of low-frequency neural activity, with encoding and retrieval linked to opposing theta phases (∼4–7 Hz), and distractor interference during maintenance linked to alpha phase (∼8–10 Hz). These findings indicate that distinct WM operations are temporally coordinated within oscillatory cycles, providing evidence that low-frequency neural activity both coordinates and segregates cognitive processes over time.

## Introduction

Growing evidence suggests that cognitive processes have functionally important rhythmic components. In the context of selective attention, behavioral studies have shown that sampling of the external environment fluctuates rhythmically at low frequencies, particularly in the theta-frequency band (3–8 Hz; Fiebelkorn et al., 2013; Galas et al., 2023; Landau & Fries, 2012; Song et al., 2014; VanRullen et al., 2007). Converging evidence indicates that these fluctuations in behavioral performance are linked to the phase of low-frequency neural oscillations across multiple brain regions involved in directing selective attention (Busch & VanRullen, 2010; Dugué & VanRullen, 2017; Fiebelkorn et al., 2018, 2019; Gaillard et al., 2020; Helfrich et al., 2018; Kienitz et al., 2018; Landau et al., 2015; Redding et al., 2026; Senoussi et al., 2025). Different low-frequency phases are associated with different spectral patterns of neural activity (Fiebelkorn et al., 2018, 2019), different functionally defined cell types (Fiebelkorn et al., 2018), different between-region interactions (Fiebelkorn et al., 2019), and different behavioral outcomes (Busch & VanRullen, 2010; Landau et al., 2015; Fiebelkorn et al., 2018; Helfrich et al., 2018; Redding et al., 2026). Together, these findings suggest that low-frequency neural oscillations may help temporally coordinate distinct, potentially competing cognitive functions. For example, rhythmic fluctuations in behavioral performance and neural activity during the deployment of selective attention may reflect the coordination of alternating brain states associated with either sampling at the current attentional focus or an increased likelihood of shifting that focus (Fiebelkorn & Kastner, 2019).

Research on working memory has similarly linked fluctuations in behavioral performance to the phase of low-frequency neural oscillations. Theta phase (∼3–8 Hz), in particular, has been associated with the maintenance and retrieval of to-be-remembered items (Abdalaziz et al., 2023; Bahramisharif et al., 2018; Cavanah & Fiebelkorn, 2026; Chota et al., 2022; Fuentemilla et al., 2010; Han et al., 2026; Kamiński et al., 2020; Peters et al., 2021; Pomper & Ansorge, 2021; Siegel et al., 2009; ter Wal et al., 2021). In some cases, different oscillatory phases have been linked to different to-be-remembered items (Abdalaziz et al., 2023; Bahramisharif et al., 2018; Kamiński et al., 2020; Siegel et al., 2009), suggesting that internal representations of different items may be strengthened during different phases within an oscillatory cycle (Lisman & Idiart, 1995; Lisman & Jensen, 2013). Such findings collectively indicate that low-frequency phase influences the accessibility of information held in working memory (i.e., during maintenance and retrieval). It remains unclear, however, whether low-frequency neural oscillations help coordinate working memory more broadly—a process that depends on multiple interconnected operations, including encoding, maintenance, retrieval, and the protection of internal representations from distraction (Baddeley, 2003; Postle, 2006). Here, we used EEG in humans to test whether low-frequency phase, measured immediately prior to key task events, was linked to behavioral performance and event-related neural responses across multiple operations of working memory.

In contrast to previous research examining whether different oscillatory phases are associated with different to-be-remembered items—that is, whether the strength of internal representations peaks at different phases for different items (Abdalaziz et al., 2023; Bahramisharif et al., 2018; Cavanah & Fiebelkorn, 2026; Chota et al., 2022; Fuentemilla et al., 2010; Han et al., 2026; Kamiński et al., 2020; Peters et al., 2021; Pomper & Ansorge, 2021; Siegel et al., 2009; ter Wal et al., 2021)—we tested whether low-frequency phase helps coordinate distinct operations of working memory. This question is conceptually related to the Separate Phases of Encoding and Retrieval (SPEAR) model, an influential account of hippocampal and cortical function during episodic memory (Hasselmo, 2025; Hasselmo et al., 2002). According to this framework, encoding and retrieval are supported during different theta phases, reducing interference between incoming and stored information. Although SPEAR was developed in the context of episodic memory (Biba et al., 2026; Kerrén et al., 2022; Rizzuto et al., 2006), it provides a useful conceptual framework for asking whether low-frequency phase temporally coordinates distinct operations of working memory (i.e., encoding and retrieval).

Based on previous findings, we further predicted that the specific low-frequency band associated with encoding and retrieval would differ from that associated with the protection of internal representations from external distractors. Whereas theta-band activity has been repeatedly linked to various aspects of working memory (Cavanagh & Frank, 2014; Herweg et al., 2020), alpha-band activity (∼9–13 Hz) has been repeatedly linked to the suppression of sensory processing (Bonnefond & Jensen, 2025; Foxe & Snyder, 2011; Jensen & Mazaheri, 2010), with some research specifically implicating alpha-band activity in the goal-directed suppression of distracting information (Redding et al., 2026; Wöstmann et al., 2019; Zhao et al., 2023; Zoest et al., 2021; Bonnefond & Jensen, 2012; but see also Redding & Fiebelkorn, 2024; van Diepen & Mazaheri, 2017; Antonov et al., 2020; Foster & Awh, 2019). The present experiment tested whether such low-frequency neural activity bridges multiple operations of working memory, influencing whether behaviorally important information is effectively encoded, successfully retrieved, and protected from distraction.

## Materials and Methods

The study protocol was approved by the Research Subjects Review Board at the University of Rochester. In accordance with the Declaration of Helsinki, each of the participants gave written informed consent prior to data collection.

### Participants

Twenty-six individuals (age range = 18–35 years; 12 females) participated in the experiment. Participants were paid $15 per hour. All participants reported normal or corrected-to-normal vision and no history of neurological conditions. Three participants were excluded from the analyses due to excessive eye movements or movement-related artifacts in the EEG (leaving an average of only 122 valid trials per excluded participant).

### Setup

The experiment was conducted on a PC equipped with a 1920 × 1080 pixel LED monitor (refresh rate: 480 Hz), in a dimly lit room. Stimuli were presented and behavioral responses were collected using MATLAB R2024b (MathWorks, Natick, MA) with the Psychophysics Toolbox extension. A chin rest was used to maintain a viewing distance of approximately 56 cm.

Gaze position was recorded monocularly at a sampling rate of 1000 Hz using an EyeLink 1000 Plus eye tracker (SR Research, Ottawa, Canada). Electroencephalographic (EEG) signals were acquired at a sampling rate of 2048 Hz using a 128-channel ActiveTwo BioSemi system (Amsterdam, The Netherlands). Electrodes were positioned according to the extended 10–10 system. Two electrooculography (EOG) electrodes were placed below and lateral to the left eye to further monitor for eye movements and blinks.

### Task and procedure

**Figure 1A** illustrates the experimental task. Each trial began when participants pressed the space bar (i.e., trials were self-paced). A black fixation dot (0.25° diameter) was then presented at the center of the screen and remained there throughout the trial. After 500 ms, a gray disk was briefly flashed at fixation for 100 ms, as a temporal cue to begin monitoring for the occurrence of a to-be-remembered item (i.e., a memory item). Following a variable delay of 750–1250 ms, this memory item, a grating stimulus (1 cycle per degree), was briefly presented at fixation for 35 ms. The grating orientation was randomly selected from 1° to 180° on each trial, and participants were instructed to remember the orientation. The memory item was followed by another variable delay of 750–1250 ms. For half of the trials, an additional grating, with a randomly chosen orientation, was then presented at fixation for 35 ms, serving as a task-irrelevant distractor. Regardless of whether a distractor was presented, there was a further variable delay of 750–1250 ms. For all trials, participants therefore had to remember the orientation of the memory item for a total of 1535–2500 ms (i.e., during the combined memory delay, with or without a distractor). At the end of each trial, a black ring (or probe) appeared at fixation to cue memory retrieval. Participants reproduced the remembered orientation by moving the mouse left or right (Ding et al., 2024; Gresch et al., 2024; van Ede et al., 2020). During this response adjustment, a randomly oriented grating appeared at fixation and rotated in accordance with the participant’s mouse movement. The response was submitted either when participants clicked the left mouse button or automatically, 4 s after participants initiated the mouse movement. All flashes, gratings, and probe stimuli had a diameter that subtended 8° of the visual angle. Memory performance, defined as the angular difference between the remembered and reported orientations, was converted into an accuracy score from 0 to 100 (0° error = 100%, 90° error = 0%) and provided to participants as feedback. Participants were instructed to respond as accurately and as quickly as possible, while maintaining fixation throughout each trial (i.e., until feedback was presented). Trials were discarded if mouse movements occurred before probe onset. Each participant completed nine blocks of 80 trials. The first block served as practice and was excluded from further analyses (i.e., leaving 640 total trials).

**Figure 1.**
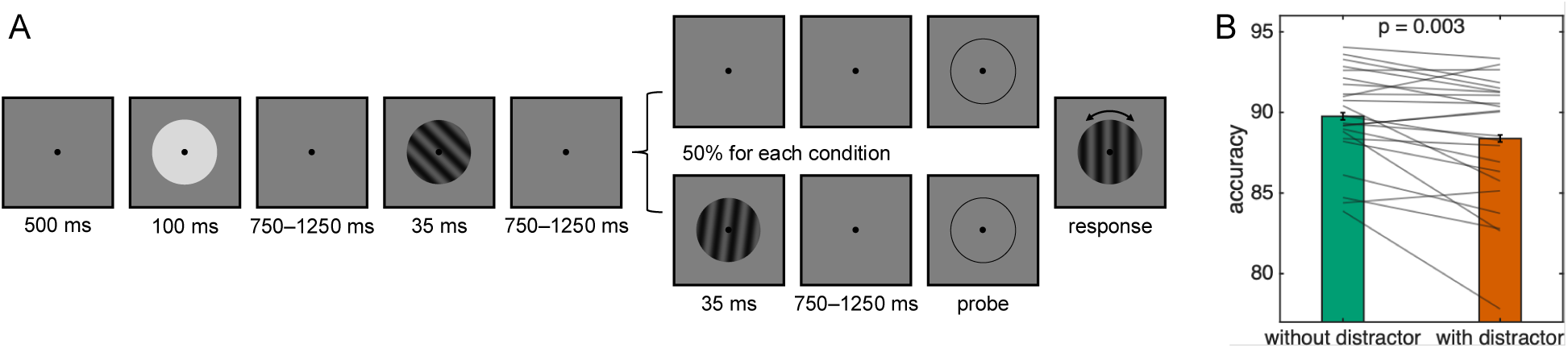
Visual working memory task and behavior results. A. Each trial began with central fixation, followed by a briefly flashed disk (i.e., a temporal cue) and a to-be-remembered grating stimulus (i.e., a memory item). For 50% of trials, a task-irrelevant distractor (i.e., another grating stimulus with a randomized orientation) appeared prior to the memory probe. Following the memory probe, participants reproduced the remembered orientation by using a mouse to rotate a randomly oriented grating. B. Memory accuracy was lower on trials with a distractor. Gray lines indicate the change across conditions for each participant.

### Analysis

Memory accuracy, across participants (n = 23), was compared for trials with and without a distractor (i.e., to assess distractor effects). For both trial types, response times shorter than 100 ms and response times longer than 5 standard deviations from the median were removed from further analyses. For this behavioral analysis, statistical significance was tested with a paired t-test. Unless otherwise indicated, we used p < 0.05 as a threshold for statistical significance.

Eye-tracking data were loaded into MATLAB using edfmex software (SR Research, Ontario, Canada) to measure participants’ gaze position and blinks. Trials in which blinks occurred, or gaze position deviated more than 3° from fixation were excluded from further analyses. As an additional step for removing trials with eye-related artifacts, the difference between the near-eye electrodes (i.e., EOG) were visually inspected to determine a participant-specific voltage threshold for further artifact rejection.

EEG data were preprocessed and analyzed in MATLAB with FieldTrip (Oostenveld et al., 2011). During preprocessing, data were epoched into three different trial events: (i) from 750 ms before to 750 ms after memory item onset (i.e., time-locked to encoding), (ii) from 750 ms before to 500 ms after probe onset (i.e., time-locked to retrieval), and (iii) from before 750 ms to 750 ms after distractor onset (i.e., time-locked to distractor processing). Epoched data were downsampled to 512 Hz and re-referenced to the average of all EEG channels (i.e., we used an average reference). The EEG data were zero-padded to 5 s, linearly detrended, and demeaned. Line noise at 60, 120, and 180 Hz was removed using a discrete Fourier transform (DFT) filter. In addition to removing trials with blinks and eye movements, a participant-specific voltage threshold, based on visual inspection, was used to identify other noise transients. If noise transients were identified on fewer than 15% of the 128 scalp EEG electrodes, data for the individual electrodes with noise transients were interpolated using the nearest four electrodes. If the number of electrodes with noise transients was greater than 15% (i.e., greater than 19 electrodes), the trial was removed from further analyses. This artifact rejection was performed using data from 750 ms before memory item onset to 500 ms after probe onset. Following the rejection of trials based on blinks, eye movements, and other noise transients, an average of 332 trials remained for further analyses. Following the approach of previous studies (Abdalaziz et al., 2023; Cavanah & Fiebelkorn, 2026; Redding et al., 2026), the current analysis focused on examining the relationship between frequency-specific phase and behavioral performance (i.e., memory accuracy). Phase prior to each event was estimated using frequency-specific Morlet wavelets. That is, Morlet wavelets were used to measure oscillatory phase just prior to encoding (prior to onset of the memory item), just prior to distractor processing (prior to onset of the distractor), and just prior to retrieval (prior to the memory probe). The number of cycles varied with frequency, using 2 cycles between 3 and 8 Hz and increasing logarithmically from 2 to 5 cycles between 9 and 55 Hz. To achieve maximal temporal precision in estimating event-related phase, wavelets were fit separately for each frequency such that the final time point of the phase estimate occurred immediately prior to the onset of the event of interest (e.g., memory item onset). This approach of prevents contamination of the phase estimates from visual responses associated with the task events. After measuring pre-event phases, we tested whether memory accuracy fluctuated as a function of oscillatory phase, separately for working memory operation (i.e., encoding, retrieval, and distractor processing). To calculate phase-accuracy relationships for each electrode-frequency combination, trials were binned according to pre-event phase using a sliding window of 180°. Behavioral performance (i.e., memory accuracy) was then calculated within overlapping phase bins. The 180° phase window was shifted from 1° to 360° in steps of 10°, thereby sampling the full oscillatory cycle. To reduce between-participant variability, behavioral measurements were normalized within each participant by subtracting the mean and dividing by the standard deviation of the behavioral measurements (see Fig 5A). To quantify the strength of the phase-accuracy relationship for each electrode-frequency combination, phase-accuracy functions (i.e., behavioral performance as a function of oscillatory phase) were first averaged across participants. A discrete Fourier transform (DFT) was then applied to each function, and the amplitude of the second Fourier component—corresponding to a one-cycle sinusoid—was extracted. This approach assumes that the ‘good’ and ‘bad’ phases for memory performance are separated by approximately 180°. The amplitude of the second Fourier component served as the metric for capturing the strength of phase-accuracy relationships (Abdalaziz et al., 2023; Fiebelkorn et al., 2013; Redding et al., 2026). See the section below for how we tested for statistical significance, while controlling for multiple comparisons across different electrode-frequency combinations.

To further examine how phases associated with either ‘good’ or ‘bad’ memory performance were related to electrophysiological measures of visual and cognitive processing, event-related potentials (ERPs) were calculated after binning trials based on the ‘good’ and ‘bad’ pre-event phases (i.e., based on the ‘good’ and ‘bad’ pre-event phases established from the phase-accuracy analyses). The frequency-electrode combination showing the strongest phase-accuracy relationship was selected to define the phases for ERP binning. The ‘good’ and ‘bad’ phase bins were defined as 120°, centered on the peak and trough of the phase-accuracy function, respectively (see Fig. 5A). Because averaging the raw EEG epochs after binning by phase can introduce phase-aligned fluctuations that complicate baseline correction, broadband filtered ERPs (1–55 Hz) were instead calculated (Redding et al., 2026). That is, prior to averaging, we applied a Hilbert transform to the broadband filtered signal (1–55 Hz) and took the absolute value to measure non-phase-aligned, broadband power. Like for the other analyses, broadband ERPs were calculated separately for different conditions (i.e., for distractor trials and no-distractor trials) and epochs of interest (i.e., time-locked to the memory item, the probe, or the distractor). Electrodes for broadband ERP analyses were selected based on the grand-average scalp topographies following memory item onset (see Fig. 5B).

To test whether the scalp topography of the phase–accuracy relationship differed between conditions, we performed a topographic analysis of variance (TANOVA) test (Murray et al., 2008). For each participant and condition, the strength of phase-accuracy functions was z-normalized across channels to form a normalized scalp map. Topographic differences were then quantified using global dissimilarity (DISS), defined as the root mean square difference between condition-averaged normalized maps. Statistical significance was assessed using a within-subject permutation test (1000 permutations), in which condition labels were randomly shuffled at the participant level and DISS was recomputed on each permutation. The p value was defined as the proportion of permuted DISS values that exceeded the observed DISS.

### Statistical Evaluation

Cluster-based permutation tests were applied to define the statistical significance for both assessing phase-accuracy relationships and ERP comparison across conditions (Cavanah & Fiebelkorn, 2026; Maris & Oostenveld, 2007; Redding et al., 2026).

For phase-accuracy relationships, the amplitude of the one-cycle sine fit to the phase-accuracy function (e.g., Fig. 5A) was used to quantify the strength of the relationship for each electrode-frequency combination. Null distributions were generated by randomly shuffling pre-event phases across trials and recalculating the phase-accuracy relationships for each participant, after which values were averaged across participants (1,000 iterations), thereby breaking any relationship between oscillatory phase and the measure of interest. The observed relationship strength was then compared against the null distribution to identify electrode-frequency combinations exceeding the 95th percentile of null values. Significant combinations were grouped into clusters across adjacent electrodes and frequencies. Cluster statistics were computed for each cluster, and for each permutation the maximum cluster statistic was retained to generate a null distribution of cluster statistics. Observed clusters with statistics exceeding the 95th percentile of this null distribution were considered statistically significant.

ERP differences between conditions (good- versus bad-phase trials) were also assessed using cluster-based permutation tests, following the same procedure described above. Statistical comparisons were performed at each time point, and clusters of contiguous time points exceeding an initial threshold of p < 0.05 were identified.

## Results

Participants reproduced the orientation of a to-be-remembered visual grating, either with or without an intervening distractor presented during the memory delay (i.e., after the memory item and before the memory probe). As shown in Figure 1B, the presence of a distractor reduced memory accuracy (*t*(22) = 3.23, *P* = 0.004), indicating that the distractor interfered with the maintenance of the internal representation (i.e., the neural representation of the to-be-remembered visual grating).

After establishing these behavioral effects, we tested whether frequency-specific phase (at 3–55 Hz) influenced visual and cognitive processes at different operations of a working memory task: (i) encoding of the memory item (i.e., the to-be-remembered visual grating), (ii) retrieval of the neural representation associated with the memory item, and (iii) visual processing of the distractor (i.e., on trials with a distractor). For each operation, we tested for a relationship between frequency-specific phase and both memory accuracy and event-related neural responses.

To calculate the relationship between frequency-specific phase and memory accuracy, we binned trials based on the phase occurring just prior to each task operation (i.e., encoding, retrieval, and distractor processing) and measured memory accuracy within each phase bin, yielding memory accuracy as a function of oscillatory phase (see Fig. 5A for an example of the phase-accuracy relationship for a single electrode-frequency combination). Assuming that the phase associated with the best memory accuracy (i.e., the “good” phase) was approximately 180° from the phase associated with the worst memory accuracy (i.e., the “bad” phase), we fit these functions with a one-cycle sine wave and used the amplitude of the fitted sine wave to estimate the strength of the phase-accuracy relationship for each electrode-frequency combination (Cavanah & Fiebelkorn, 2026; Fiebelkorn et al., 2013; Redding et al., 2026).

Figure 2 shows the strength of the relationships between memory accuracy and frequency-specific phase measured just prior to encoding (i.e., prior to memory item onset; Fig. 2A–F) and retrieval (i.e., prior to probe onset; Fig. 2G–L), for both trials without distractors and trials with distractors. Non-significant electrode-frequency combinations are set to zero. Here, we controlled for multiple comparisons by using non-parametric cluster-based permutations (Cavanah & Fiebelkorn, 2026; Maris & Oostenveld, 2007; Redding et al., 2026). Averaging phase-accuracy relationships across electrodes (Fig. 2B) revealed that phasic modulation of memory encoding in lower-frequency bands was stronger when a distractor was subsequently presented during the memory delay, peaking within the theta (5–6 Hz) and alpha (8–12 Hz) bands (Fig. 2D). In comparison, phase-accuracy relationships were weaker or absent during trials without a subsequent distractor (Fig. 2A and C). Figures 2E and F show the scalp topographies and significant electrode clusters associated with peaks in phase-accuracy relationships during trials with a distractor. For theta-band activity (Fig. 2E), stronger phase-accuracy relationships over occipital regions are consistent with previous work linking theta phase to the retrieval operation of working memory tasks (Cavanah & Fiebelkorn, 2026). The present results demonstrate that the phase of lower-frequency neural activity also influences the encoding operation of a working memory task—with effects being strongest within the theta band. These phase-accuracy relationships during the encoding operation were apparent despite frequency-specific phase being measured up to seconds before the subsequent memory probe and behavioral response. Relationships between pre-encoding phase and memory accuracy, however, were largely dependent on the occurrence of an intervening distractor.

**Figure 2.**
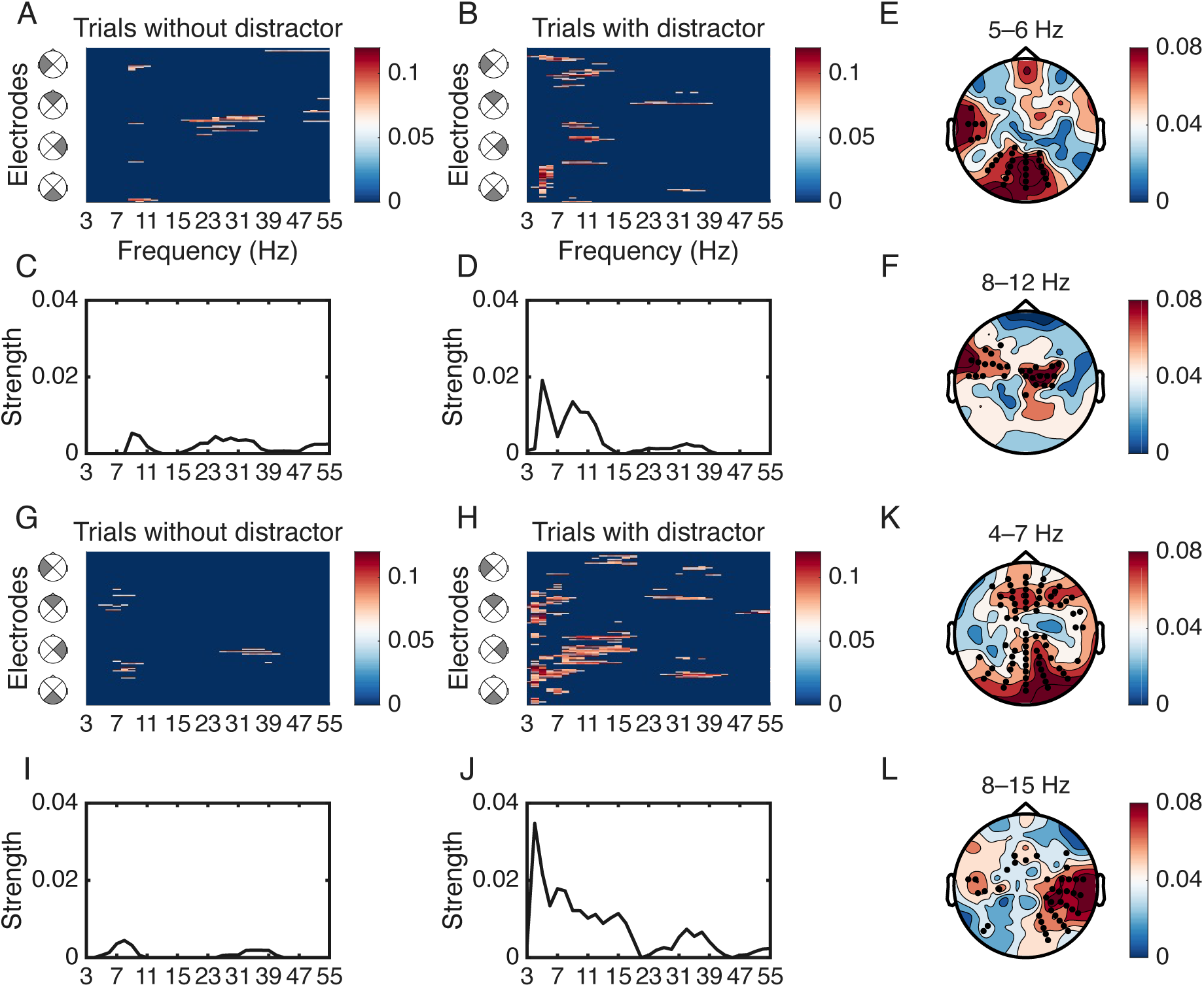
Phase–accuracy relationships associated with memory item and probe onset. (A–F) Significant phase-accuracy relationships associated with memory item onset (non-significant values have been zeroed). (A, B) Frequency-electrode combinations showing significant phase-accuracy relationships in trials either without (A) or with (B) a distractor. (C, D) Average phase-accuracy relationships across electrodes in trials either without (C) or with (D) a distractor. (E, F) Scalp topographies illustrating the spatial distribution of phase-accuracy relationships in the theta (5–6 Hz) and alpha (8–12 Hz) bands for trials with a distractor. (G–L) Same analyses as in (A–F), but for phase-accuracy relationships associated with probe onset. Black dots on the topographic plots indicate significant electrodes.

Using the same procedures that linked pre-encoding phase with memory accuracy, we tested for relationships between frequency-specific phase and other processing operations of the working memory task. Previous work has reported a relationship between frequency-specific phase and the retrieval of to-be-remembered information (Abdalaziz et al., 2023; Cavanah & Fiebelkorn, 2026; Zheng et al., 2024). The results shown in Figure 2G–L are largely consistent with those findings, demonstrating statistically significant relationships between lower-frequency phase just prior to the memory probe and memory accuracy (p < 0.05, after correction for multiple comparisons), with the strongest relationships in the theta band (at 4–7 Hz, Fig. 2K) and a secondary peak in the alpha band (at 8–15 Hz, Fig. 2L). Similar to relationships between pre-encoding phase and memory accuracy (Fig. 2A–F), relationships between pre-retrieval phase and memory accuracy were (i) strongest at frontal and occipital electrodes (Fig. 2K, L) and (ii) almost exclusively observed on trials with distractors (see Discussion).

Memory accuracy was most strongly influenced by the phase of theta-band activity, measured both prior to the memory item (i.e., prior to encoding) and prior to the memory probe (i.e., prior to retrieval). To test whether the scalp distributions of the phase-accuracy relationships differed between these two working memory operations, we performed TANOVA separately at each theta-band frequency (4–8 Hz). No significant topographic differences were observed at any tested frequency (DISSs = 0.25–0.30, all p > 0.45), indicating that the spatial configurations of the phase-accuracy relationships were broadly similar across the two operations, consistent with shared neural generators. Theta-band activity in various brain regions has been linked to the temporal coordination of cognitive processes. For example, different theta phases in the network that directs selective attention are associated with either (i) attention-related sampling of behaviorally important information from the external environment or (ii) the shifting of attentional resources across the external environment (Fiebelkorn et al., 2013; Fiebelkorn & Kastner, 2019; Landau & Fries, 2012; VanRullen, 2016). Similarly, a prominent computational model of episodic memory proposes separate theta phases for encoding and retrieval (i.e., the SPEAR model; Hasselmo et al., 2002; Hasselmo, 2025). We therefore tested whether the *pre-encoding phase* associated with better memory accuracy (i.e., the “good” phase) differed from the *pre-retrieval phase* associated with better memory accuracy. The significant clusters of phase-accuracy relationships associated with encoding and retrieval included 16 shared electrode-frequency combinations within the theta band (5–7 Hz). Figure 3A shows an example of the phase-accuracy functions for one of these shared electrode-frequency combinations, and Figure 3B shows an associated polar histogram of the “good” phases from each participant, separately for the encoding and retrieval operations. Circular Watson-Williams tests demonstrated that these phases differed significantly (p < 0.05) for 15 of the 16 shared electrode-frequency combinations (Fig. 3C), with an average phase difference of 134°. The present findings are therefore consistent with the SPEAR model, indicating that different theta phases promote encoding versus retrieval. Although not perfectly anti-phasic, the theta phase associated with better encoding is generally associated with worse retrieval, and vice versa.

**Figure 3.**
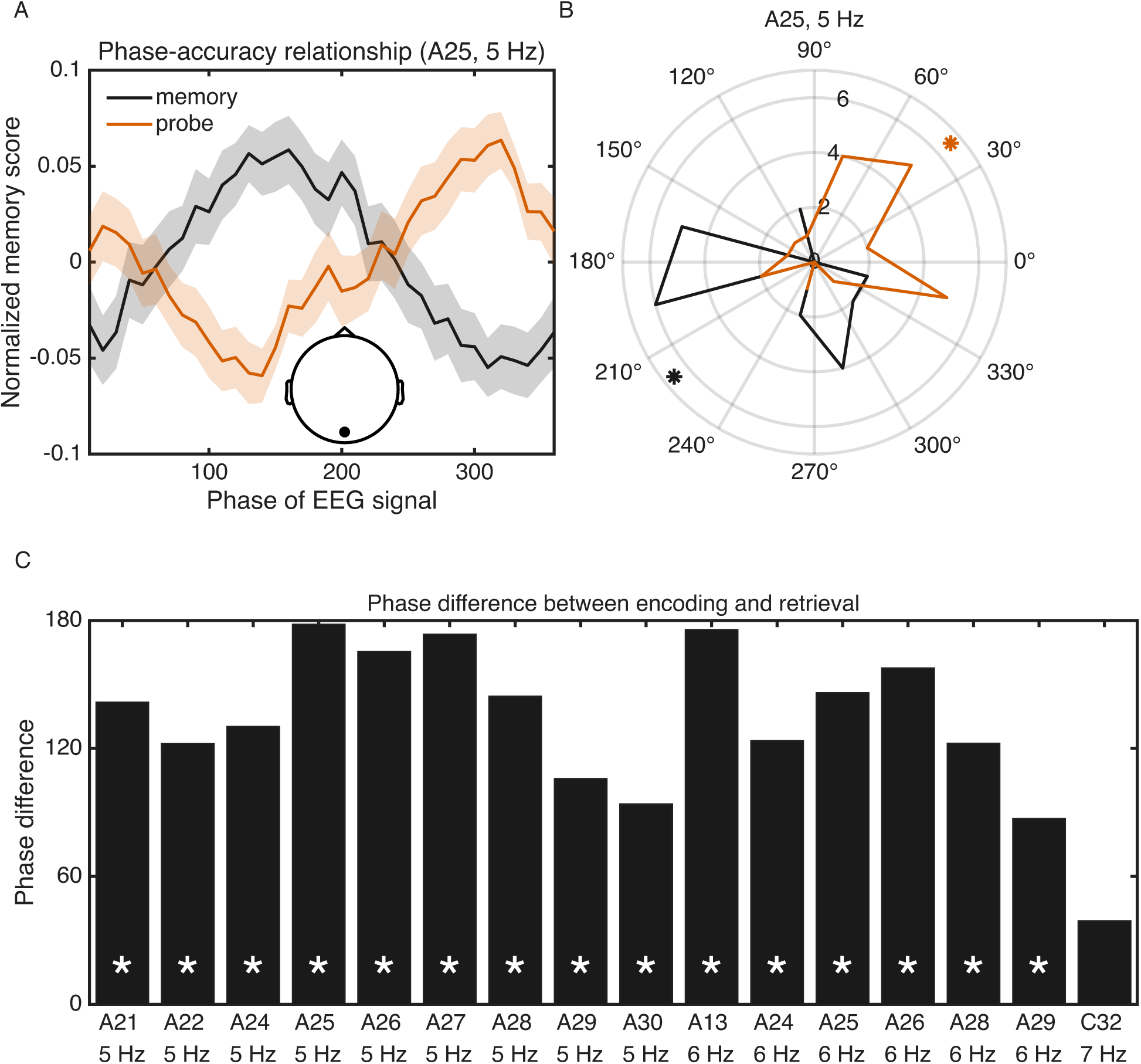
Comparison of the “good” theta phases associated with significant phase-accuracy relationships for both memory item and probe onset. (A) Phase-accuracy relationships with phase measured just prior to memory item onset (black line) or just prior to probe onset (red line), specifically at electrode A25 and 5 Hz. (B) Circular histograms of the peak “good” phases (i.e., the phases associated with better memory accuracy) shown in (A), plotting the phase of the fitted one-cycle sine waves for each participant’s phase-accuracy function (n = 23), separately for memory item (black) and probe onset (red). Angular mean phases for each working memory operation are indicated by black and red stars. (C) Phase differences between memory item- and probe-associated “good” phases for each significant frequency-electrode combination (16 total electrodes). White stars indicate statistically significant phase differences (p < 0.05).

Previous research has shown that lower-frequency phase, specifically alpha phase (∼9–13 Hz), can influence distractibility during visual-target detection (i.e., during selective processing of the external environment) (Bonnefond & Jensen, 2012; Redding et al., 2026; Solís-Vivanco et al., 2018; VanRullen, 2016; Wöstmann et al., 2020; Zhao et al., 2023; Zoest et al., 2021). Here, we similarly tested whether lower-frequency phase influenced distractor processing. Figure 4 shows the relationship between pre-distractor phase and memory accuracy (p < 0.05, after correction for multiple comparisons). We again observed a statistically significant relationship between lower-frequency phase and memory accuracy, with the strongest effects in the alpha range (8–10 Hz; Fig. 4D), consistent with previous findings. In comparison, both the encoding and retrieval operations were associated with stronger phase-accuracy relationships in the theta range (Fig. 2).

**Figure 4.**
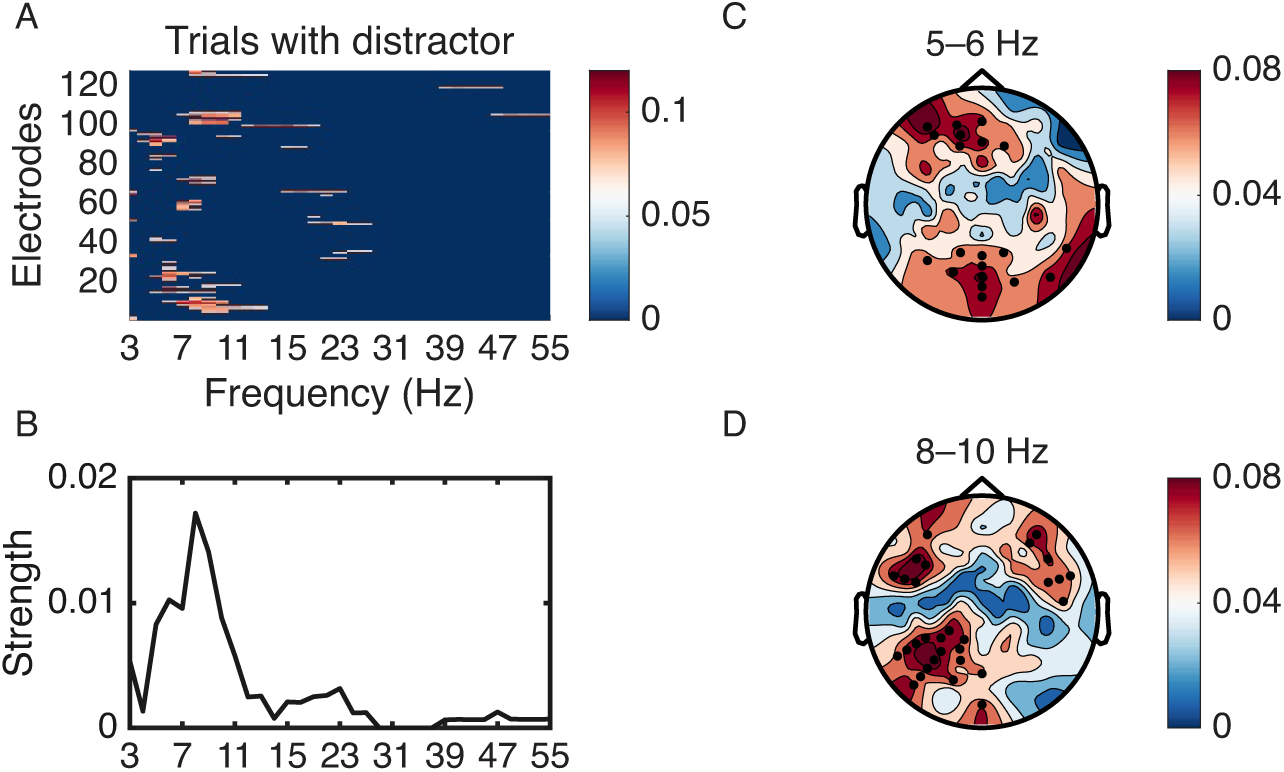
Phase-accuracy relationships associated with distractor onset. (A) Significant phase-accuracy relationships (non-significant values have been zeroed). (B) Average strength of the phase-accuracy relationships across electrodes. (C, D) Scalp topographies showing peak phase-accuracy modulation in the theta- (5–6 Hz; C) and alpha-frequency (8–10 Hz; D) bands.

To further investigate how lower-frequency phase influences cognitive processes at different operations of a working memory task, we tested whether lower-frequency phase was also related to neural activity associated with memory item onset (i.e., encoding), memory probe onset (i.e., retrieval), or distractor onset (i.e., distractor processing). Here, we first focused on the relationship between pre-encoding phase and the neural response to memory items. We split trials with distractors (i.e., trials where we previously observed strong phase-accuracy relationships) into two bins based on the pre-encoding phases associated with either better or worse memory accuracy: a bin centered on the “good” phase (±60°) and a bin centered on the “bad” phase (±60°). We specifically used the “good” and “bad” phases from the frequencies and electrodes with the strongest phase-accuracy relationships (i.e., Electrode A24 at 5 Hz; Fig. 5A). We then generated broadband event-related potentials (ERPs) time-locked to memory item onset (Redding et al., 2026), separately for each phase bin. ERP analyses focused on posterior electrodes showing the strongest early visual response to the memory item (100–200 ms following onset; Fig. 5B). As determined with a cluster-based permutation approach (see Methods), the “good” theta phase was associated with a significantly stronger neural response (p < 0.05, after correction for multiple comparisons) than the “bad” theta phase, starting approximately 160 ms after memory item onset, specifically on the trials with a subsequent distractor. These results confirm that the pre-encoding phase of theta-band activity (i.e., the theta phase just prior to memory item onset) influenced processing of the memory item—in addition to a subsequent influence on memory accuracy (Fig. 5C). The scalp topography of the difference wave was again consistent with occipital generators—similar to the previously observed phase-accuracy relationships (Fig. 2E)—with the strongest effects at occipital electrodes (Fig. 5D). Because we also observed a significant, though weaker, relationship between pre-encoding alpha phase and memory accuracy (i.e., at 9 Hz; Fig. 2B), we further tested for a relationship between pre-encoding alpha phase and broadband ERPs, time-locked to memory item onset (Supplemental Fig. 1); however, that analysis revealed no phase-related difference. That is, the relationship between pre-encoding phase and memory-item-evoked neural responses was limited to the theta band.

**Figure 5.**
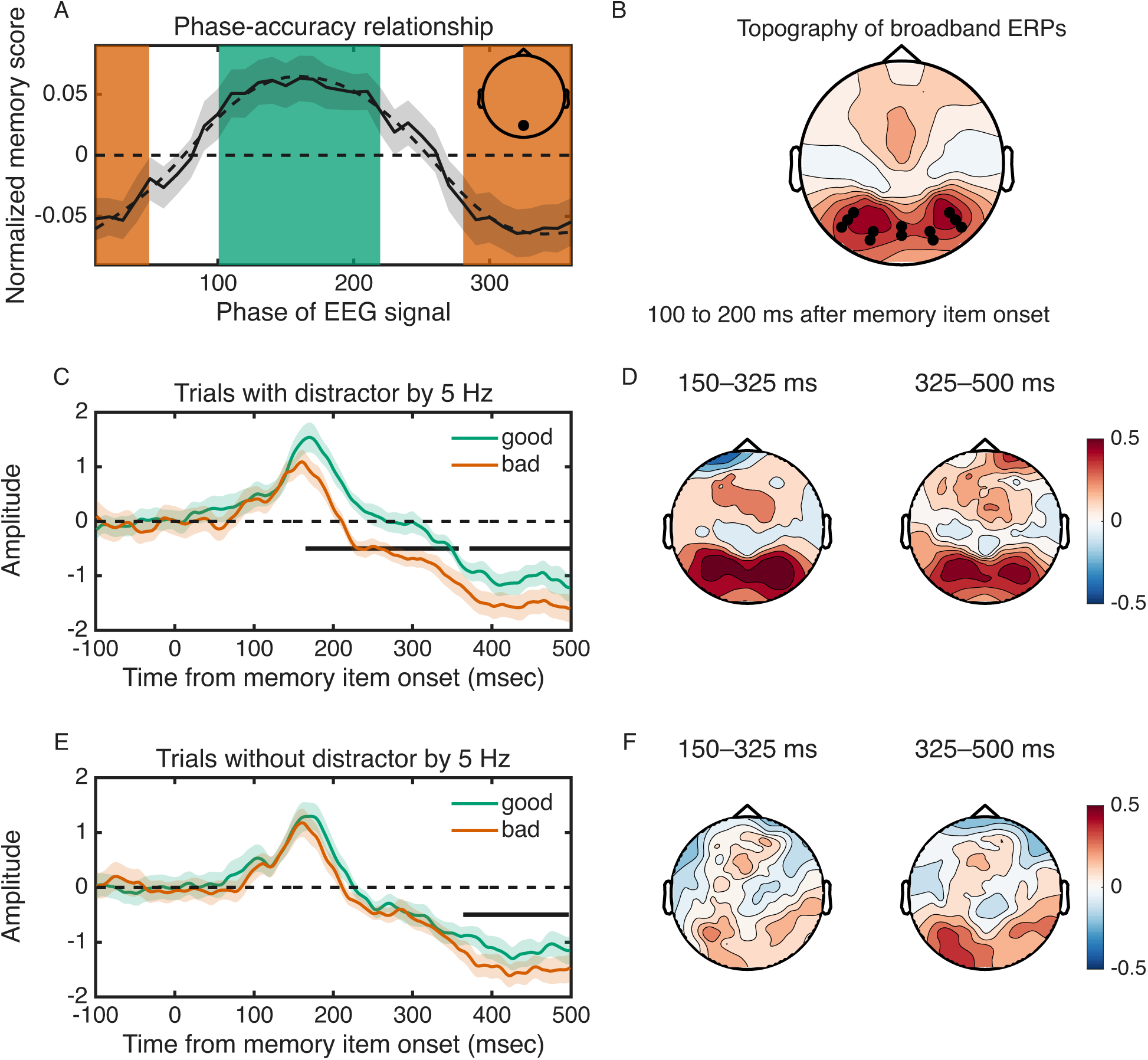
Broadband ERP differences associated with pre-encoding theta phase and memory item onset. (A) Example phase-accuracy function, showing the original function (solid black line) and a fitted sine wave (dashed black line). The amplitude of the fitted sine wave was used as the strength of the phase-accuracy relationship. Phases within ±60° of the peak were classified as good phases (green), whereas phases within ±60° of the trough were classified as bad phases (red). (B) Scalp distribution of memory-item-induced broadband ERPs (at the peak amplitude, from 100–200 ms poststimulus), with posterior electrodes used for subsequent analyses indicated by black dots. (C) Broadband ERPs binned by 5-Hz phase (at electrode A24, which had the strongest phase-accuracy relationship), aligned to memory item onset for trials with a distractor. (D) Topographic broadband ERP difference corresponding to (C). (E) Broadband ERPs binned by the same electrode and frequency as in (C) aligned to memory item onset for trials without a distractor. (F) Topographic broadband ERP difference corresponding to (E). Horizontal black lines indicate the time windows with significant differences in panels (C) and (E).

Given that pre-encoding theta phase influenced memory-item evoked responses on trials with distractors, the same should be true for trials without distractors—there was no difference between the two trial types prior to the occurrence of the distractor (i.e., when the memory item was presented). We therefore also measured the relationship between pre-encoding theta phase and memory-item evoked responses on trials without distractors. Note that for these trials, there was no relationship between pre-encoding theta phase and memory accuracy (Fig. 2A). To bin trials based on pre-encoding theta phase, we used the “good” and “bad” phases from the trials with distractors, specifically from the electrode with the strongest phase-accuracy relationship on those trials (Fig. 5A). Figure 5E shows the results of this analysis. Despite basing the binning procedure on the results from a different set of trials (i.e., from trials with distractors), we again observed a significant, phase-dependent difference between the broadband ERPs time-locked to memory item onset (p < 0.05, after correction for multiple comparisons). Although the difference did not reach statistical significance until approximately 360 ms after memory item onset, the general pattern of results was similar to that from trials with distractors (Fig. 5C), including the scalp topographies of the difference wave (Fig. 5F). These results further confirm that the pre-encoding phase of theta-band activity (i.e., the theta phase just prior to memory item onset) influences processing of the memory item, resulting in stronger or weaker encoding.

Finally, we used the same approach to test (i) whether pre-retrieval phase influenced broadband ERPs associated with memory probe onset and (ii) whether pre-distractor phase influenced broadband ERPs associated with distractor onset. Using the electrode-frequency combinations with the strongest phase-accuracy relationships—separately for retrieval and distractor processing—to split trials into “good” and “bad” phase bins revealed no significant differences between broadband ERPs, time-locked either to memory probe onset or to distractor onset (Supplemental Fig. 2).

## Discussion

Here, we examined the relationship between frequency-specific phase and behavioral performance across multiple working memory operations. For encoding, retrieval, and distractor processing, low-frequency phase—measured immediately before each task event—predicted memory accuracy. Encoding and retrieval were most strongly linked to behavioral performance through theta phase (at ∼4–7 Hz), with similar scalp topographies, consistent with shared neural generators. Distractor processing, by contrast, was most strongly linked to behavioral performance through alpha phase (at ∼8–10 Hz). These frequency-specific findings are broadly consistent with prior research demonstrating both theta-band links to working memory (Cavanagh & Frank, 2014; Herweg et al., 2020) and alpha-band links to sensory suppression (Bonnefond & Jensen, 2025; Foxe & Snyder, 2011; Jensen & Mazaheri, 2010). The present results extend this work by showing how frequency-specific neural activity spans—and may help coordinate—multiple working memory operations.

Previous research has linked theta-band activity to multiple aspects of working memory, including causal modulation of working memory performance (Albouy et al., 2017; Hoy et al., 2016; Lee & D’Esposito, 2012; Polanía et al., 2012; Riddle et al., 2020; Violante et al., 2017), increased working memory load (Gevins et al., 1997; Jensen & Tesche, 2002; Raghavachari et al., 2001; Sauseng et al., 2010), and phase-specific representations of to-be-remembered items during maintenance and retrieval (Abdalaziz et al., 2023; Bahramisharif et al., 2018; Han et al., 2026; Kamiński et al., 2020; Siegel et al., 2009). The present results confirm that theta phase measured just prior to retrieval is associated with fluctuations in memory accuracy, perhaps reflecting theta-dependent fluctuations in the strength of memory representations (i.e., internal representations of the to-be-remembered items; Bahramisharif et al., 2018; Fuentemilla et al., 2010; Han et al., 2026; Kamiński et al., 2020; Siegel et al., 2009). These findings support previous research linking theta phase to fluctuations in the coding of to-be-remembered items. For example, Bahramisharif et al. (2018) demonstrated that neural representations of sequentially presented, to-be-remembered items were associated with distinct theta phases. We have previously hypothesized that such theta-dependent fluctuations during working memory maintenance and retrieval may reflect temporal coordination of transient bursts of neural activity (Lundqvist et al., 2016) that refresh short-term synaptic changes (Abdalaziz et al., 2023). This would be consistent with “activity-silent” theories of working memory (Kamiński & Rutishauser, 2020; Lundqvist et al., 2018; Miller et al., 2018; Stokes, 2015), which propose that synaptic changes contribute to short-term storage of to-be-remembered items alongside persistent neural activity (Constantinidis et al., 2018).

In addition to confirming a relationship between theta phase and retrieval, the present results demonstrate that pre-encoding theta phase was similarly associated with fluctuations in memory accuracy. Notably, this behaviorally relevant neural activity was measured up to seconds before the behavioral response. Pre-encoding theta phase modulated not only memory accuracy but also the neural response to memory item onset: a stronger broadband response was observed when the memory item occurred during the “good” theta phase (i.e., the pre-encoding theta phase associated with better memory accuracy) relative to during the “bad” theta phase, consistent with stronger encoding. Importantly, the present experimental task included a variable memory delay, meaning that the pre-encoding phase was non-predictive of the pre-retrieval phase. The present results therefore demonstrate that theta phase independently influenced these two operations of working memory (i.e., encoding and retrieval). Classical models emphasize stimulus strength, sensory gain, and attentional resource allocation as key determinants of successful encoding (Baddeley et al., 1993; Carrasco & Barbot, 2019; Luria et al., 2016; Oberauer, 2019). The present results indicate that these factors operate within constraints imposed by oscillatory neural dynamics. Even when stimulus properties and task demands are held constant, the neural system cycles through theta-dependent phases of enhanced and diminished receptivity to new information—such that encoding depends, in part, on alignment between stimulus onset (e.g., memory item onset) and endogenous oscillatory states.

Previous research suggests that theta phase influences retrieval through fluctuations in the strength of internal representations (Abdalaziz et al., 2023; Bahramisharif et al., 2018; Han et al., 2026; Kamiński et al., 2020), but how does theta phase influence encoding? One possibility is that theta phase influences encoding through fluctuations in attention-related sampling. Selective processing of external stimuli is characterized by theta-rhythmic fluctuations in attention-related neural and behavioral effects (Fiebelkorn & Kastner, 2019; Fries, 2023; Re et al., 2023; Schroeder et al., 2010), and it is well-established that attention and working memory encoding are interdependent (Awh et al., 2006; Chun & Turk-Browne, 2007; Gazzaley & Nobre, 2012; Postle, 2006). Moreover, there appears to be a shared theta-rhythmic process for selective sampling of both external stimuli and internal representations (i.e., information maintained in working memory), with a consistent phase for enhancing behaviorally important information regardless of source (Cavanah & Fiebelkorn, 2026). This raises the possibility that theta-rhythmic fluctuations during encoding and retrieval reflect a shared sampling process that operates across external and internal information.

However, other aspects of the present findings suggest that this interpretation is incorrect, or at least incomplete. First, the “good” phase for encoding (i.e., the phase associated with better memory accuracy) significantly differed from the “good” phase for retrieval. This contrasts with our recent finding of a shared “good” phase for attention-related sampling of both external stimuli and internal representations (Cavanah & Fiebelkorn, 2026). Second, pre-encoding phase influenced the neural response to memory item onset, whereas pre-retrieval phase did not influence the neural response to probe onset. This further indicates that theta-band activity interacted differently with encoding- and retrieval-related neural processes—and was not simply phasically boosting behaviorally important external information prior to encoding and internal information prior to retrieval, via a shared attention-related “sampling” phase (Cavanah & Fiebelkorn, 2026; Fiebelkorn & Kastner, 2019).

The present results are instead compatible with an influential model of hippocampal and cortical function during episodic memory that predicts distinct theta phases for encoding and retrieval (Hasselmo, 2025; Hasselmo et al., 2002). Similar to other theories addressing how frequency-specific neural activity temporally coordinates competing functions or representations (Fiebelkorn & Kastner, 2019; Lisman & Jensen, 2013), the SPEAR model proposes that separate theta phases—within a single oscillatory cycle—are associated with either encoding or retrieval, with each phase being linked to unique patterns of neural activity (Hasselmo, 2025). This alternation between functional states may help prevent interference between incoming and currently maintained information. While evidence for separate encoding and retrieval phases is well established in rodents (Hasselmo, 2025), evidence in humans has been relatively limited (but see Biba et al., 2026; Kerrén et al., 2022; Rizzuto et al., 2006). The present results provide evidence in humans that distinct pre-encoding and pre-retrieval theta phases are associated with better memory performance during a working memory task.

For both encoding and retrieval, phase-accuracy relationships were stronger on trials with distractors. Because phase-dependent modulation of behavioral performance requires behavioral variability, high overall memory accuracy may have reduced the observable strength of phase-accuracy relationships, particularly on trials without a distractor. Alternatively, the distractor itself may have influenced the neural activity underlying phase-accuracy relationships by interacting with latent frequency-specific dynamics (Jung et al., 2025), thereby strengthening or re-introducing theta-rhythmic fluctuations of internal representations. Consistent with this possibility, previous work has shown that task-irrelevant stimuli presented during a memory delay can transiently re-engage neural processes linked to memory maintenance (Sprague et al., 2016; Stokes et al., 2013; Wolff et al., 2017). As a post hoc test of this idea, we split trials by median memory delay and re-examined the relationship between pre-retrieval phase and memory accuracy separately for trials with and without distractors. Despite reduced trial counts, this analysis suggests that phase-accuracy relationships weakened over longer delays on trials without a distractor but persisted on trials with a distractor (Supplemental Figure 3). That is, coupling between low-frequency phase and memory accuracy diminished over time in the absence of a distractor but was preserved when a distractor was present, perhaps because the distractor re-engaged theta-rhythmic processes that strengthened—or re-strengthened—internal representations.

In addition to this indirect influence on theta-related phase-accuracy relationships during encoding and retrieval, the direct influence of the distractor on memory accuracy was modulated by pre-distractor alpha phase, consistent with previous results and alpha’s proposed role in sensory suppression (Bonnefond & Jensen, 2025; Foxe & Snyder, 2011). While this relationship was consistent with gated inhibition of distractor processing (Jensen & Mazaheri, 2010), alpha phase did not influence the distractor-evoked neural response (see Redding et al., 2026). This pattern of results suggests that alpha phase modulated the susceptibility of internal representations to distractor interference rather than sensory processing of the distractor itself.

In conclusion, the present results reveal a frequency-specific neural architecture that helps coordinate multiple working memory operations, thereby influencing when information can be most effectively encoded, retrieved, and protected from distraction. Rather than being uniform over time, working memory performance depends on the alignment between task events and alternating neural states that promote different working memory operations—from encoding through retrieval. Distractor interference both depends on frequency-specific neural activity (i.e., pre-distractor alpha phase) and modulates relationships between frequency-specific neural activity and other working memory operations (e.g., the relationship between theta phase and retrieval). Together, these findings suggest that low-frequency neural oscillations provide a temporal framework that bridges and perhaps coordinates the multiple operations of working memory.

## Acknowledgements

This work was supported by grants from the National Science Foundation (NSF 2120539) and the Searle Scholars Program to I.C.F, and from the National Institutes of Health (NEI T32EY007125 and NIMH F31MH141973) to P.C. We would like to thank Hourieh Hayati and Jordan Klembczyk for their help with data collection.

## Supplemental information

**Supplemental Figure 1.**
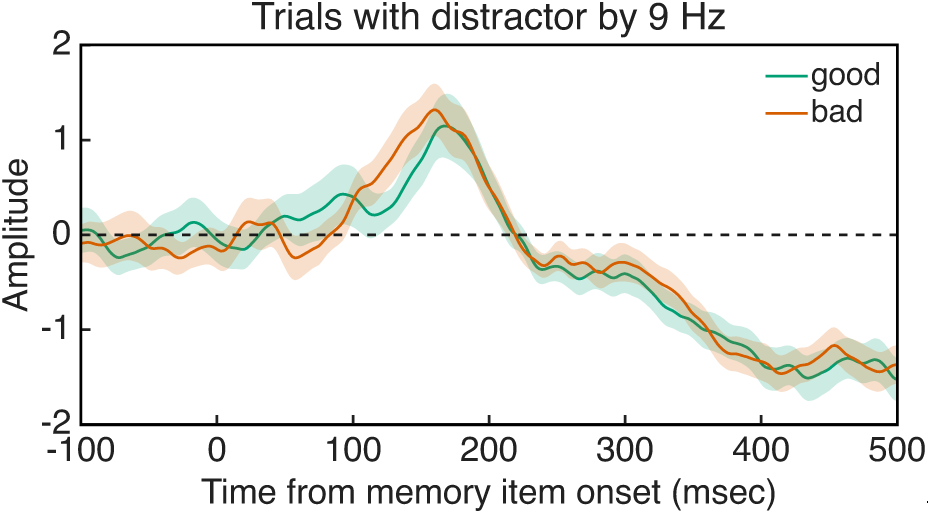
Alpha-band phase did not modulate broadband ERPs time-locked to memory item onset in trials with a distractor. Trials were binned according to the 9-Hz phase at electrode D8 (i.e., the electrode with the strongest phase-accuracy relationship in the alpha-frequency band) into good-phase (green) and bad-phase (red) trials.

**Supplemental Figure 2.**
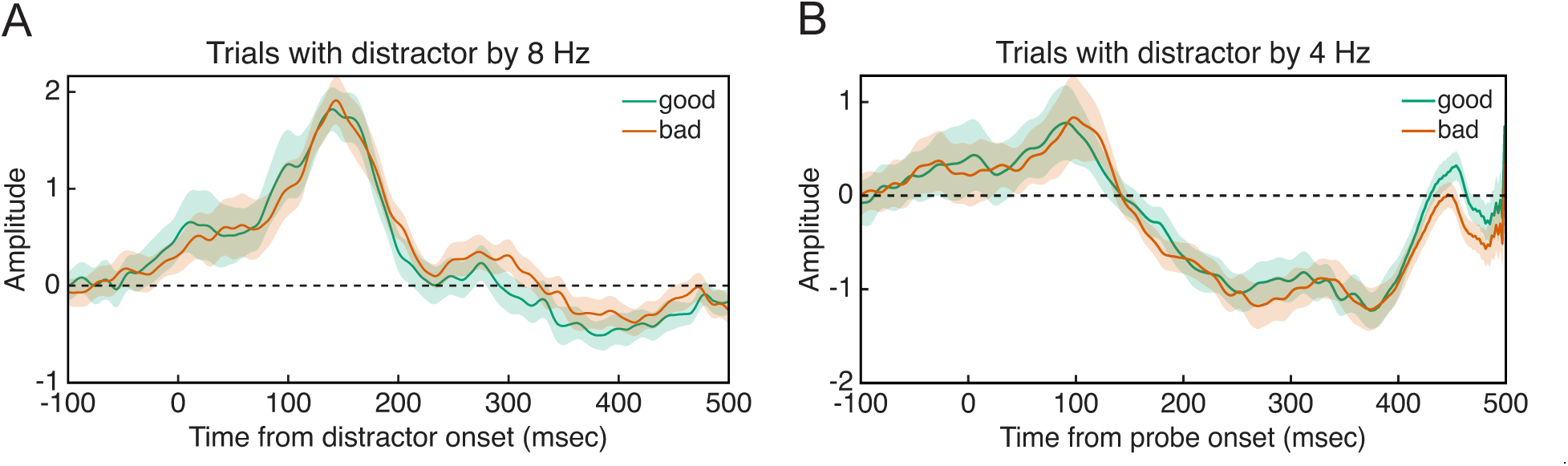
Broadband ERPs associated with phase-dependent trials. (A) ERPs time-locked to distractor onset, with trials binned according to the 8 Hz phase at electrode A9 into good-phase (green) and bad-phase (red) trials. (B) ERPs time-locked to probe onset, with distractor trials binned according to the 4 Hz phase at electrode A32 into good-phase (green) and bad-phase (red) trials.

**Supplemental Figure 3.**
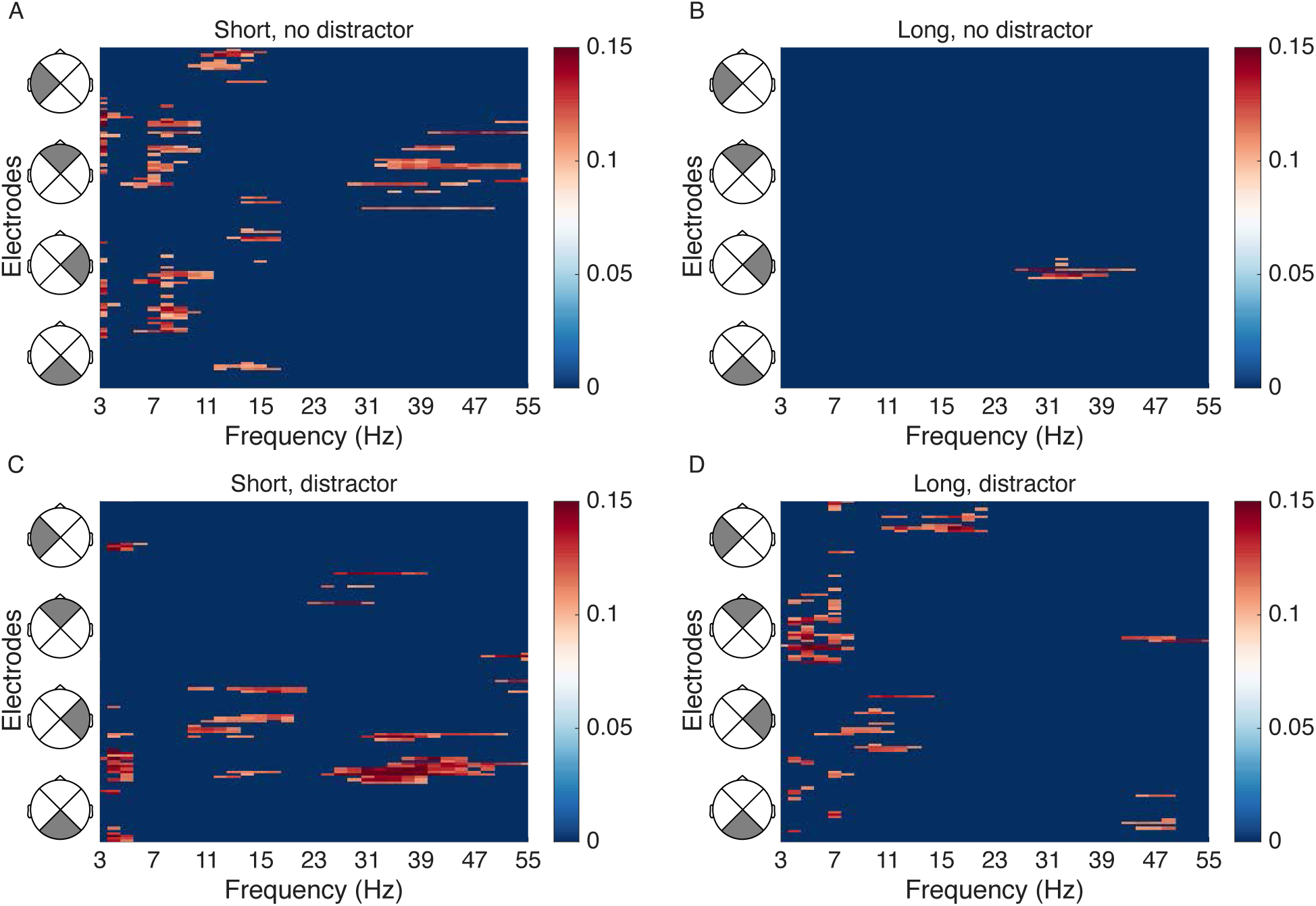
Phase-accuracy relationships associated with probe onset. Trials were grouped by memory delay (short vs. long) and distractor presence. (A–D) Significant phase-accuracy relationships across frequency–electrode pairs, with non-significant values set to zero. (A, B) No-distractor trials for short and long delays, respectively. (C, D) Distractor-present trials for short and long delays, respectively.

